# Different transcriptional responses by the CRISPRa system in distinct types of heterochromatin in *Drosophila melanogaster*

**DOI:** 10.1101/2022.02.21.481378

**Authors:** Andrea Ortega-Yáñez, Samantha Cruz-Ruiz, Martha Vázquez, Mario Zurita

## Abstract

Transcription factors (TFs) activate gene expression by binding to elements close to promoters or enhancers. Some TFs can bind to heterochromatic regions to initiate gene activation, suggesting that if a TF is able to bind to any type of heterochromatin, it can activate transcription. To investigate this possibility, we used the CRISPRa system based on dCas9-VPR as an artificial TF in *Drosophila*. dCas9-VPR was targeted to the *TAHRE* subtelomeric element, an example of constitutive heterochromatin, and to promoters and enhancers of the HOX *Ultrabithorax* (*Ubx*) and *Sex Combs Reduced* (*Scr*) genes in the context of facultative heterochromatin. dCas9-VPR robustly activated *TAHRE* transcription, showing that although this element is heterochromatic, dCas9-VPR was sufficient to activate its expression. In the case of HOX gene promoters, although these genes are epigenetically silenced by Polycomb complexes, both were ectopically activated. When the artificial TF was directed to enhancers, we found that the expression pattern was different compared to the effect on the promoters. In the case of the *Scr* upstream enhancer, dCas9-VPR activated the gene ectopically but with less expressivity; however, ectopic activation also occurred in different cells. In the case of the *bxI* enhancer located in the third intron of *Ubx*, the presence of dCas9-VPR is capable of increasing transcription initiation while simultaneously blocking transcription elongation, generating a lack of functional phenotype. Our results show that transcription can be activated in any type of heterochromatin by CRISPRa; nevertheless, its effect on transcription is subject to the intrinsic characteristics of each gene or regulatory element.

**Significance:** Whether transcription only depends on activating factors binding to chromatin, even though it is found in a silent state as heterochromatin, remains an open question. In this work, we addressed this question using the CRISPRa system via dCas9-VPR as a synthetic transcriptional activator in *Drosophila*. This activator was directed to a constitutive heterochromatin element and to promoters and enhancers of two HOX genes, which in the tissues where they are not expressed, are present as facultative heterochromatin. In all cases, the CRISPRa system was able to activate transcription, showing that its sole presence is sufficient for this to occur. Although transcription in constitutive heterochromatin was very robust, in the case of promoters and enhancers of HOX genes, the degree of expressivity, penetrance and ectopic effect was different between promoters and enhancers. These results indicate that the presence of a synthetic activator can activate transcription by binding to transcriptional regulatory elements; however, its effect depends on the particular characteristics of each one. These results show how artificial transcription factors can be used to understand transcription regulation at the organismal level.

## Introduction

Gene expression mediated by RNA polymerase II (RNPII) in eukaryotic cells is modulated by the action of activators that recruit the basal transcription machinery to the promoter to form the preinitiation complex (PIC)(1,2). In some cases, such as the transcriptional activation mediated by nuclear hormone receptors, the GAGA factor and the heat shock factor, the transcriptional activator binding site could be located near the promoter(3–5). However, in metazoans, most of the developmentally regulated genes, as well as genes that respond to different signal transduction pathways, are activated by the action of enhancers (6,7). These cis-regulatory elements (CREs) are recognized by multiple transcription factors (TFs) and are able to operate at large distances from the promoter by the formation of chromatin loops through the interaction between transcription factors and components of PIC(8–10). In general, the interaction of the TF with coactivators and/or with elements of the PIC occurs through a specific region known as the activation domain(11). Transcriptional activators recruit protein complexes that modify and/or remodel chromatin to maintain transcriptional permissive regions known as euchromatin(12–14).

On the other hand, in nontranscribed regions, the chromatin conformation, called heterochromatin, is highly compacted into a “closed stage”(15). In general, two types of heterochromatin have been identified based on the degree of compaction, which is linked to specific histone modifications. Facultative heterochromatin is present in regions that include silenced genes in specific cell types, and its promoters are enriched with the repressive mark tri-methylation of lysine 27 of histone 3 (H3K27me3)(16,17). This histone modification is introduced by components of the Polycomb group of genes in *Drosophila*, which include Polycomb Repression complexes 1 and 2 (PRC1, PRC2) through the action of the methyl transferase Enhancer of Zeste *[E(z*)](18). The action of PRC1 and PRC2 ensures the epigenetic silencing of genes during development(19). In contrast, constitutive heterochromatin, considered to have a higher state of compaction of chromatin, is mostly present in subtelomeric and centromeric sequences, and it is characterized by enrichment of the trimethylation of lysine 9 of the histone 3 (H3K9me3) mark(20).

It has been established that the conformation of chromatin is a determinant in the activation of gene expression. However, many TFs, mostly pioneers, recognize specific DNA elements in compact chromatin and are able to alter the structure of the nucleosome and recruit factors to activate transcription and open the chromatin(14). It seems that this is the first event in transcription activation, suggesting that sending a transcriptional activator to any region in the chromatin could potentially cause its transcriptional activation.

The CRISPR/Cas9 system has been modified either to activate (CRISPRa) or to repress (CRISPRi) transcription(21). To induce the transcription of specific genes, a dead Cas9 (dCas9) has been fused to the activation domains of several TFs, with dCas9-VPR being one of the most widely used(22). VPR is comprised of three activation domains derived from VP6, p65 and Rta TFs, and it has been used to activate transcription in a variety of models(23)^-23^. The dCas9 fused to activator domains is usually directed upstream of the transcription start site (TSS) of a specific gene, and recently, it has been used to activate CREs such as enhancers(24, 25).

Based on this information, our main question was whether the dCas9-VPR system could activate transcription in any kind of heterochromatin in a complete organism. For this purpose, we used Drosophila as a model organism since the combination of the GAL4-*UAS* system with CRISPRa-dCas9-VPR allows precise genetic manipulation in a complete animal(26). To test this system in constitutive heterochromatin, we analyzed whether the subtelomeric *TAHRE* element, which is maintained as silenced constitutive heterochromatin, can be activated by dCas9-VPR. For the activation of facultative heterochromatin, we selected the HOX genes *Ultrabithorax* (*Ubx*) and *Sex combs reduced* (*Scr*). These genes are highly regulated during fly development, and their regulated silencing is maintained by PRC1 and PRC2(27).

Intriguingly, dCas9-VPR was able to transcribe both types of heterochromatic elements during development. In the subtelomeric element, transcription was efficiently activated, showing that dCas9-VPR can act as a pioneer TF. In the case of HOX genes, dCas9-VPR was able to activate transcription; however, the effect on the activation of transcription was different between enhancers and promoters. In particular, the occupation of the synthetic transcriptional activator in the *bxI* enhancer located in a *Ubx* intron increases transcription initiation, but at the same time, it seems to block the passage of RNPII, generating a loss-of-function *Ubx* phenotype. These results show that the activation of transcription in different types of heterochromatin by the dCas9-VPR system generates distinct transcriptional responses that depend on their location and the roles they may have in the activation of transcription.

## Results

### Ectopic transcriptional activation of subtelomeric sequences in *D. melanogaster* by the CRISPRa system

In *Drosophila*, telomeres are maintained by the transposition of *non-long terminal repeat* (*non-LTR*) retrotransposons. These retro-transposable elements (*rTE*), also referred as the *HTT* array, are known as *Het-A, TART* and *TAHRE*(28). These elements are silenced via the Piwi-piRNA pathway, which requires the transcription of segments of the rTE during oogenesis to induce the formation of heterochromatin(29). We chose this region to test the transcription of constitutive heterochromatin using the CRISPRa system. We designed a double guides RNAs (dgRNAs) directed upstream of the 5’ region of the *TAHRE* element (Figure 1A, Sup Table 1). *A UAS-dCas9-VPR* transgene was expressed in all cell types and developmental stages under the control of the GAL4*-*UAS system using the *Act5C-GAL4* driver (Fig. 1A). We used the *Act5C* driver since it is not strong; therefore, the levels of dCas9-VPR are not high, similar to the native levels of most TFs.

**Fig 1.**
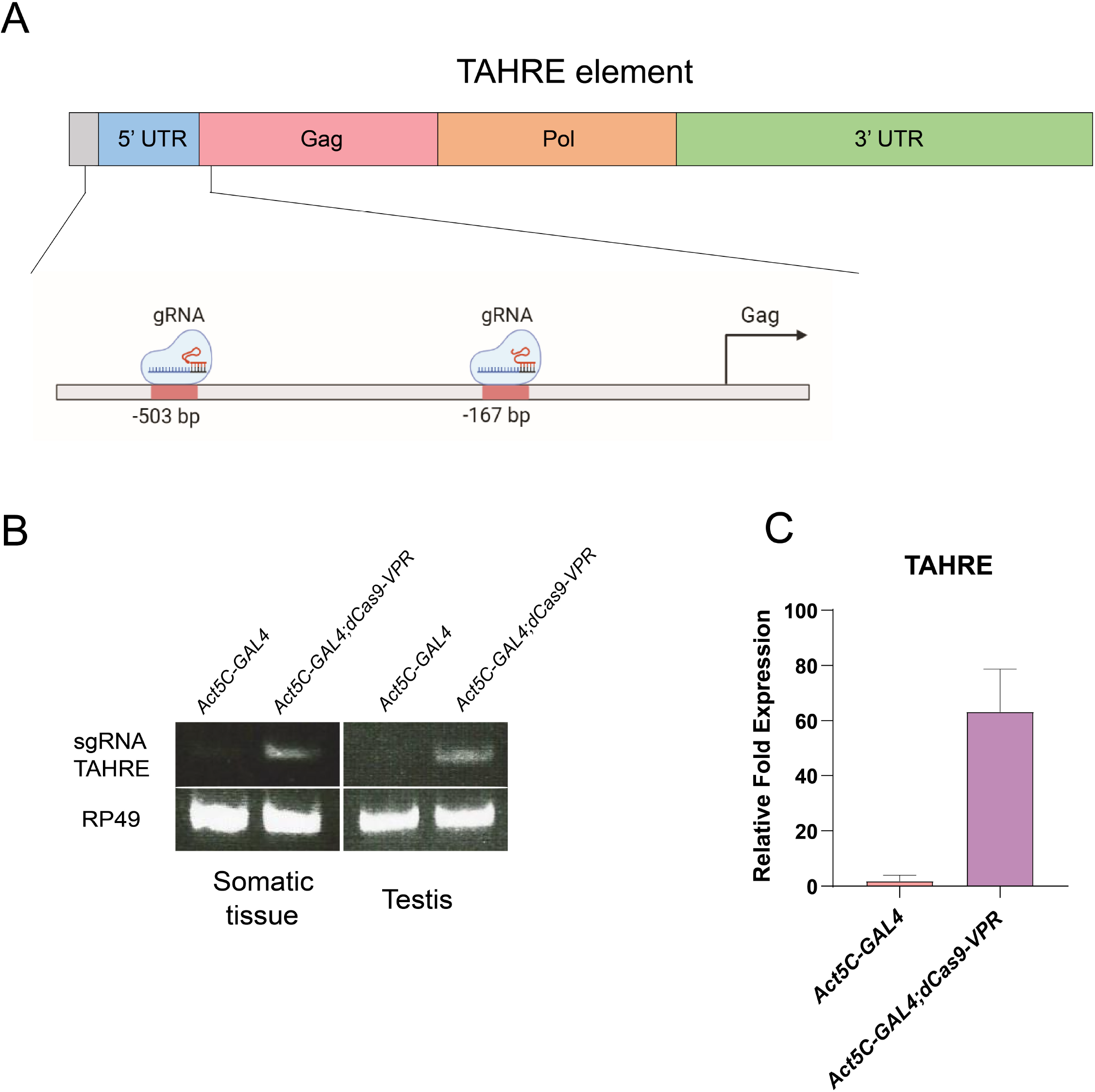
Transcription activation of the heterochromatic element *TAHRE* mediated by the dCas9-VPR. A) Representation of the *TAHRE* element present in subtelomeric regions and the location of the designed gRNAs. The position of the gRNAs are indicated in Supp. Table 1. B) Expression of *TAHRE* in somatic tissue in adult males and in testis evaluated by RT-PCR. *Act5C-GAL4;UAS-dCas9-VPR* indicates the RT-PCR from total RNA from flies in which the dCas9 is directed to the *TAHRE* 5’region compared to the *Act5C-Gal4* control flies. *rp49* mRNA was used as a loading control. C) Transcript accumulation of *TAHRE* evaluated by qRT-PCR experiments from two biological replicates. Transcript levels of Rp49 were used as a reference.

Intriguingly, we did not detect any phenotype in these flies during development. However, we analyzed the ectopic expression of the *TAHRE* element by standard semiquantitative RT–PCR and qRT–PCR in adults. We observed that in adult flies with the CRISPRa genotype, the expression of *TAHRE* was induced in both testis and somatic tissue compared with the control flies, where we did not detect any transcript (Fig. 1B). Then, by qRT–PCR, we detected an increase of approximately 60 times of the *TAHRE* transcript in comparison with the control flies (Fig. 1C). These results indicate that dCas9-VPR can direct transcription of the heterochromatic *TAHRE* element.

### Ectopic expression of the *Ubx* gene by directing the CRISPRa system to its promoter

HOX genes can be considered a typical example of facultative heterochromatin because they are epigenetically spatiotemporally silenced by PRC1 and PRC2 complexes during *Drosophila* development(18,30,31). To determine whether the CRISPRa system can activate the transcription of this type of gene in cells in which it is epigenetically silenced, we selected the HOX gene *Ubx. Ubx* controls the identity of thoracic segment 3 (T3) and the anterior part of abdominal segment 1 (A1)(32–34). During larval stages, *Ubx* is expressed in the haltere and T3 metathoracic leg (T3 leg disc) imaginal discs. In cells where *Ubx* should not be expressed(35,36), it is found as facultative heterochromatin enriched with the mark H3K27me3(37). *Ubx* is transcribed from a single promoter, and its regulatory elements are extended by approximately 100 Kb (Fig. 2A)(38).

**Fig 2.**
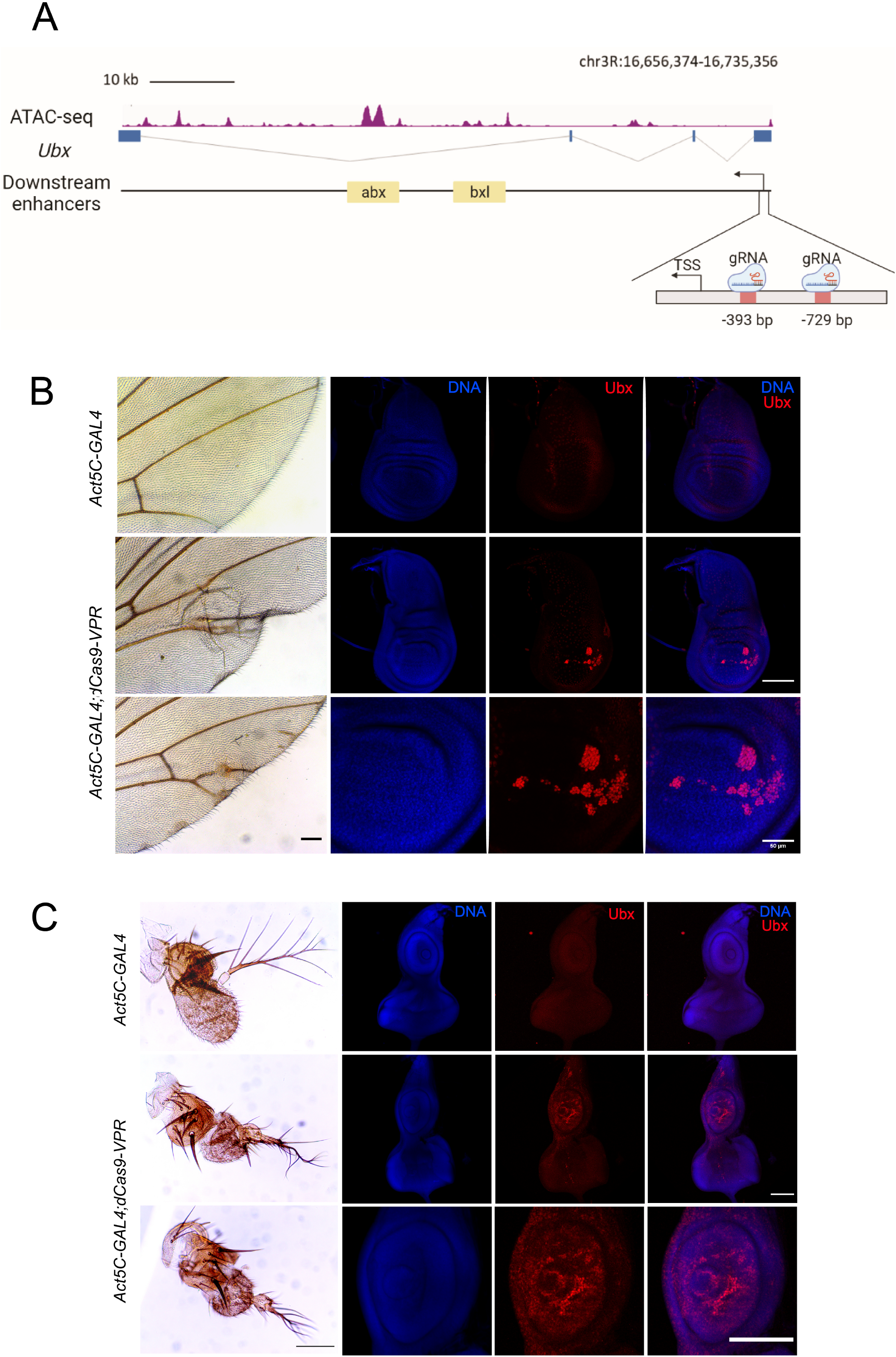
*Ubx* expression mediated by the dCas9-VPR directed to the *Ubx* promoter generates ectopic variegated expression causing homeotic transformations. A) Map of the *Ubx* gene showing its genomic organization and the position of the gRNAs designed upstream of its promoter. The enhancers localized inside the body of the gene were mapped using previously reported ATAC-seq data(48). The genome coordinates of the gRNAs positions are indicated. B) Phenotypes generated by the ectopic expression of *Ubx* in adult wings and its corresponding imaginal disc comparing with control lacking dCas9-VPR activator. In the zoom of the pouch it is shown the variegated phenotype in *Act5C-GAL4;dCas9-VPR* disc. C) Ectopic expression of *Ubx* in the eye-antenna disc and the transformed phenotypes from antennas to legs in adult organisms. The identification of Ubx was determined by immunostanings and DNA was stained with DAPI, Barr: 100 μm.

Following the previous strategy, we expressed dCas9-VPR under the control of the *Act5C* driver in all fly tissues. The gRNAs were directed upstream of the transcription initiation site of the *Ubx* gene (Fig. 2A; Sup Table 1). Interestingly, although dCas9-VPR was expressed ubiquitously, we did not detect any reduction in viability. However, phenotypes that are typical of *Ubx* ectopic expression were observed in wings and antennae with a penetrance of 63% (Sup. Table 2). For instance, the mutant phenotype in adult wings is observed predominantly in the second (2P) and third (3P) posterior regions as well as in cubital (L4) and distal (L5) veins of the wing (Fig. 2B). Additionally, adult wings presented extra cell clusters in L4 and L5 and either extra or absence of veins, as well as extra bristles and patches of cells in 2P and 3P (Fig. 2B). In addition, mutant phenotypes consistent with an initial antenna-to-leg transformation were found. We observed the appearance of bristles in the arista and in the bulge of this structure (Fig. 2C). At the base of the arista, we also observed the appearance of extra bristles protruding from antennomere 3 (A3) (Fig. 2C). These phenotypes are consistent with those described for an antenna-to-leg transformation(39) and references therein.

Following these results, we analyzed whether Ubx is expressed in dCas9-VPR wing and eye-antenna imaginal discs given that these wild-type discs do not normally express this gene (40,41). In dCas9-VPR-expressing wing discs, patches of cells expressing *Ubx* were observed to be located mostly in the pouch along the dorsal-posterior area (Fig. 2B). For the eye-antenna imaginal disc, we detected the ectopic presence of *Ubx* predominantly in the antenna region and less so in the eye section (Fig. 2C). This pattern of *Ubx* expression in the wing and in the antenna-eye imaginal discs is consistent with the mutant phenotypes observed in the corresponding structures in the dCas9-VPR adults. These data suggest that the dCas9-VPR directed to the *Ubx* promoter is able to activate the ectopic expression of this gene in wing and antenna-eye imaginal discs, resulting in variegated expression, particularly of the posterior region in the wing disc.

Since the expression of dCas9-VPR was directed by a ubiquitous driver, we wondered if the ectopic expression of *Ubx* would increase by directing the expression of dCas9-VPR to a specific region of the wing disc. For this purpose, we used the specific wing *apterous* (*ap*-GAL4) and *MS1096* drivers. As shown in Sup. Fig. 1, the effect on *Ubx* expression was similar to that observed using the actin driver. When the tubulin driver was used, which is stronger than *Act5C*, the transformation of antennae to legs was more dramatic (Sup. Fig. 1).

In summary, the dCas9-VPR system, when directed to the *Ubx* promoter, can induce its variegated ectopic expression in some but not all imaginal discs, generating changes in the cell identity that are manifested in adult organs. In addition, ectopic *Ubx* expression in the wing imaginal disc preferentially occurs in specific regions, indicating that within the disc, there are cells that seem to be more permissive to derepress *Ubx* than others. These results indicate that the presence of an artificial transcriptional activator is sufficient to activate *Ubx* despite its normal silenced state in the wing and eye-antenna discs.

### Ectopic expression from the *Scr* promoter by the CRISPR/dCas9-VPR system

To extensively analyze the activation of transcription by the CRISPR/dCas9-VPR system in genes silenced by Pc, we also evaluated the *Sex comb reduced* (*Scr*) gene. *Scr is a* HOX gene involved in the differentiation of the labial and prothoracic segments(42). Its ectopic phenotypes are identified in male adults by the presence of extra sex combs in the mesothoracic (T2) and metathoracic (T3) legs, similar to those present in the prothoracic leg (T1) in wild-type males(43). Similar to *Ubx, Scr* is highly regulated during development, with specific enhancers localized upstream and inside the second intron, directing its expression at different locations and developmental stages(42). The *Scr* gene encodes two isoforms that are transcribed from two overlapping promoters and, therefore, from two TSSs(43). Two gRNAs were designed to bind to a region between the Polycomb Response Elements (PREs) 4/9, which are binding sites for PCR1 and PRC2 that cover the two TSSs(44) (Fig. 3A; Sup. Table 1). Male adult flies expressing dCas9-VPR and gRNAs under the *Actin5C* driver had a strong fully penetrant gain-of-function *Scr* phenotype, showing the presence of ectopic sex combs in T2 and T3 legs (Fig. 3B-C). The expressivity (number of combs per leg) of the phenotype varies between different organisms (Fig. 3B), but 100% of males exhibit this homeotic transformation (Fig. 3C).

**Fig. 3.**
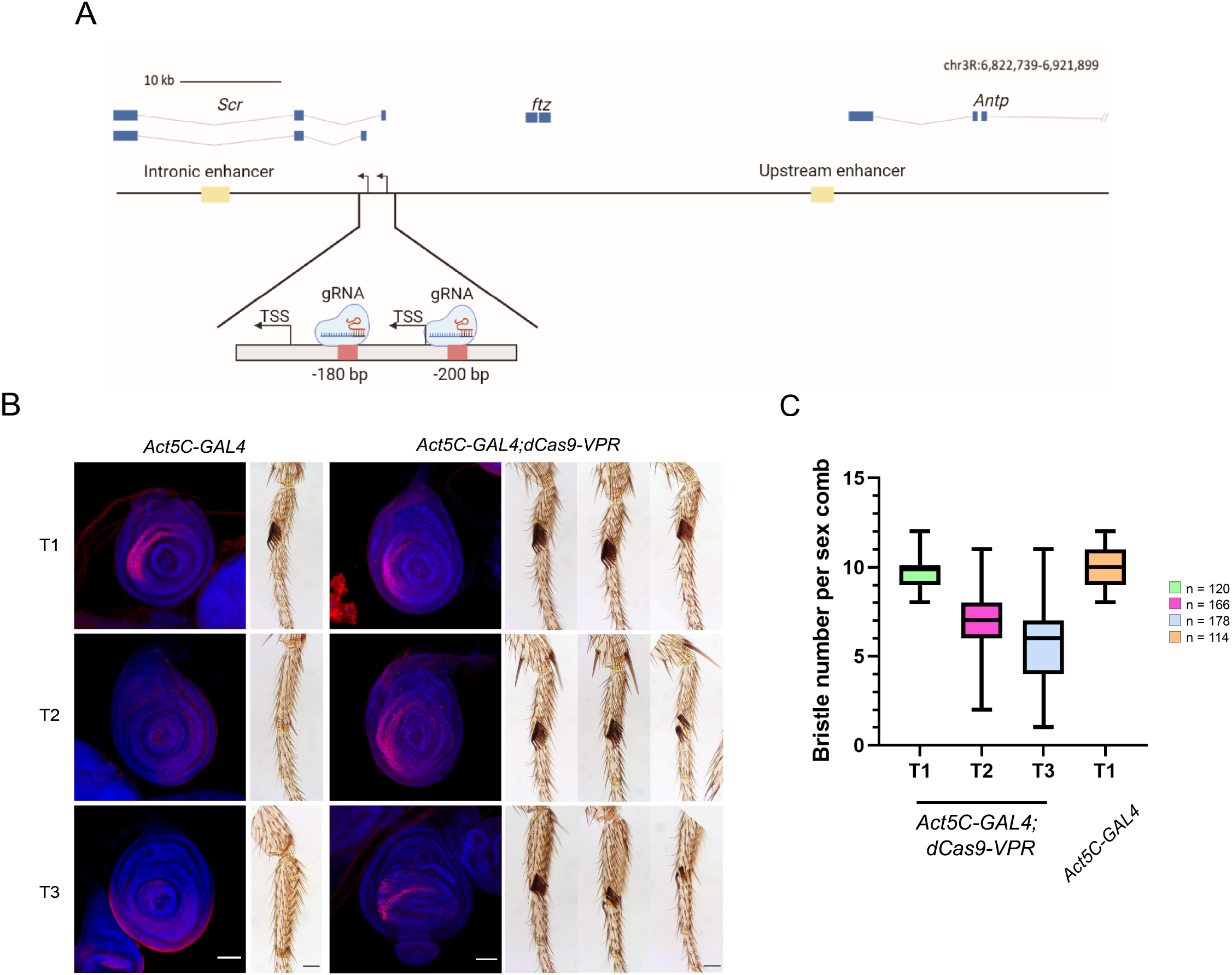
dCas9-VPR directed to the *Scr* promoter cause the transformation of the 2^nd^ and 3^rd^ pair of legs to the 1^st^ pair legs. A) Map of the *Scr* gene showing its genomic organization, the position of enhancers that regulate *Scr*, as well as the position where the gRNAs were sent upstream of the *Scr* promoters. The genome coordinates of the gRNAs position is indicated. B) Ectopic expression of *Scr* by immunostaining using the actin driver in the 2^nd^ and 3^rd^ legs discs (Barr: 50 μm.), along with the transformed phenotypes generated in adult legs. C) Quantification of the expressivity indicated by the number of extra sex combs in the legs two and three (T2, T3) compared with control *Act5C-*GAL4 flies.

Next, we analyzed the *Scr* expression pattern in leg imaginal discs in males by immunostaining (Fig. 3B). As expected, we found ectopic expression of *Scr* in the T2 and T3 leg discs. In summary, the occupancy of dCas9-VPR of the *Scr* promoter region induces its expression in other leg discs where it is normally silenced, generating a typical Polycomb mutant phenotype (31).

### Targeting the dCas9-VPR to the *bxI* enhancer abolishes *Ubx* <u>expression</u>

Recently, it has been reported that sending the CRISPRa system to enhancers can activate gene expression of the target gene(24). Thus, we analyzed the effect of directing dCas9-VPR to enhancers of the HOX genes *Ubx* and *Scr. Ubx* is regulated during development by different enhancers that are tissue-specific(45). The *anterobithorax* (*abx*) and *bithorax* (*bx*) regions contain several CREs that are located in the third intron of *Ubx*, approximately 30 kb downstream of the TSS (Fig. 4A). These regions are functional in haltere imaginal discs(46). The *bxI* region is responsible for part of the enhancer activity of *bx*(47). Recently, CREs in this region have been dissected more precisely by chromatin accessibility assays(48,49).

**Fig. 4.**
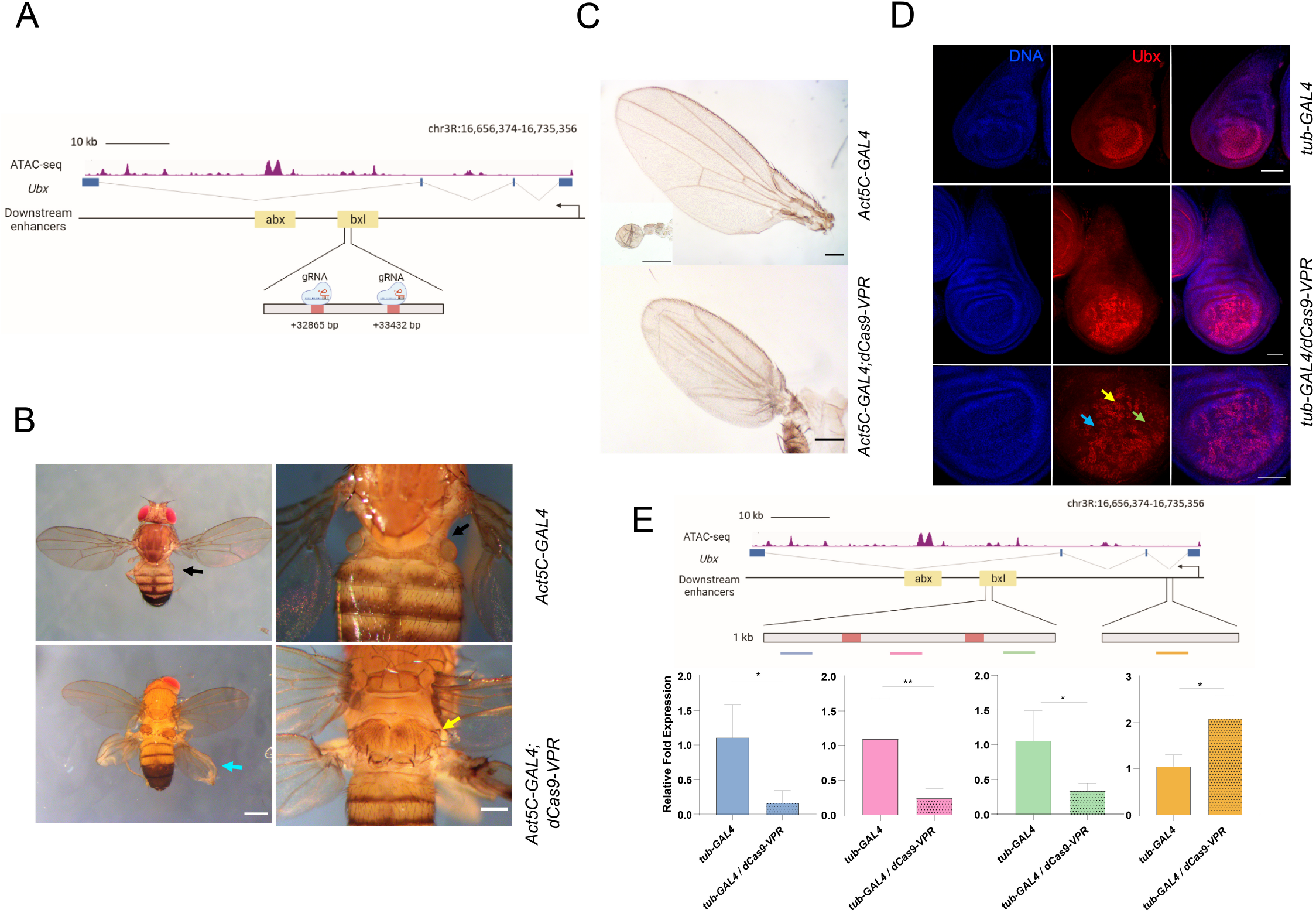
The presence of dCas9-VPR in the *bxI* enhancer generates a typical loss of function phenotype of *Ubx*. A) Genomic organization of the *Ubx* locus showing the position of the gRNAs designed inside the *bxI* enhancer localized inside the third intron of the gen; the open chromatin regions were mapped using previously published ATAC-seq data(48). The genome coordinates of the gRNAs position are indicated. B) Homeotic transformations generated by the occupancy of the dCas9-VPR in the *bxI* enhancer. The arrows indicate the location of the halters in the *Act5C-GAL4* control fly (black) and the transformation of the metanotum (yellow) and the halteres to wings in the *Act5C-GAL4;dCas9-VPR* flies (blue). C) Transformation of the halters to wings comparing the size versus the control structures. D) Immunostainings of Ubx in haltere discs when dCas9-VPR is sent to the *bxI* enhancer. The arrows mark cells that do not express Ubx (blue), express low levels of Ubx (green) and normal levels of Ubx (yellow). E) qRT-PCR of the transcripts that surround the occupancy of the dCas9-VPR in the *bxI* enhancer and of the nascent *Ubx* transcript. At least three independent experiments were performed (see material and methods). Transcript levels of Rp49 were used as a reference. The positions in the *Ubx* gene of the amplicons analyzed are indicated in the figure and its location is indicated in Sup. Table 2. *p <0.05 ; **p <0.01.

We directed the dCas9-VPR system to the *bxI* enhancer at 13,540 bp from the *abx* CRE(47,48) based on public ATAC-seq data that show this region is an open chromatin area in haltere cells(48) (Fig. 4A). Intriguingly, transgenic flies with this construct presented a homeotic haltere-to-wing transformation as well as transformation of the metanotum corresponding to the anterior part of the third thoracic segment (T3a) to exhibit characteristics of the anterior part of the second thoracic segment (T2a) (Fig. 4B), consistent with *Ubx* loss-of-function phenotypes (Fig. 4B, C). The mutant phenotype had a penetrance of 100% (Sup. Table 3). Interestingly, no other defects were evident. This phenotype is generally present in adult flies when *Ubx* expression is reduced in the T3 thoracic segment, meaning that a reduction in *Ubx* activity in the haltere disc was generated by sending the dCas9-VPR transcriptional activator to the *bxI* enhancer. We analyzed Ubx levels by immunostaining the imaginal discs. First, we noticed that haltere discs with dCas9-VPR directed to the *bxI* showed a partial to total transformation to wing disc morphology (Fig. 4D). Additionally, we observed a variegated Ubx distribution in the distal region of the disc as well as a different distribution of Ubx compared to the wt disc (Fig. 4D).

These results suggest that, contrary to what we expected, the binding of dCas9-VPR to the *bxI* enhancer generates a *Ubx* loss-of-function phenotype, although immunostaining indicates that Ubx levels and its distribution are only partially affected. This raises the question of whether the binding of dCas9-VPR in the intronic *bxI* affects enhancer activity or, as an alternative, blocks the elongation of RNPII, as has been reported when dCas9 is present downstream of the TSS of some genes(50–52). Additionally, it is also possible that *Ubx* overexpression in haltere discs may generate a similar homeotic transformation, since *Ubx* negatively regulates its own expression(36).

To determine which of these hypotheses is correct, we performed qRT–PCR experiments of the *bxI* enhancer region around the dCas9-VPR binding sites, as it is known that there is a correlation between enhancer activity and the transcription of the enhancer RNA (eRNA)(53–55). We performed RT–qPCR of the three regions around the sites that recognize the gRNAs (Fig. 4E). Clearly, there was a decrease in the transcripts arising from this region of *bxI* (Fig. 4E), indicating that the presence of dCas9-VPR reduced transcription in that region. However, this result does not differentiate whether the synthetic transcription factor affects the function of the enhancer or prevents RNAPII elongation.

To answer this question, we quantified nascent pre-*Ubx*-RNA by RT–qPCR, targeting the first intron near the *Ubx* TSS (Fig. 4E). Intriguingly, the *Ubx* nascent transcript levels increased by twofold in haltere discs in which dCas9-VPR is targeted to *bxI*. However, *Ubx* transcription was not detected in the wing disc in the same flies (Sup. Fig. 2). These results indicate that the occupancy of dCas9-VPR in this CRE enhances its effect on *Ubx* transcription initiation only in the haltere disc. However, since this enhancer is within the body of the gene, we propose that the generation of the loss-of-function phenotype is because it blocks RNPII elongation in some cells.

### dCas9-VPR induces the overexpression of *Scr* when it is directed to the upstream enhancer

The results presented up to this point with *Ubx* have shown that its effect on the induction of transcription by dCas9-VPR may not be the same when it is targeted to a promoter as it is when targeted to an enhancer. Thus, we tested the transcriptional effect of the dCas9-VPR system when directed to a nonintronic enhancer that is located far away from the gene it acted on, as is the case of the *Scr* upstream enhancer, *ScrE*(42). This enhancer is at -33 Kb with respect to the *Scr* TSS of the two promoters, between the 3’ end of *fushi tarazu* (*ftz*) and the 3’ end of *Antennapedia* (*Antp*) (Fig. 5A). *Scr*^*E*^ includes a subfragment of 439 bp that contains several putative binding sites for homeodomain TFs and that can direct the expression of *Scr* to specific regions in the T1 leg disc, including the primordia of the sensory organ(42). We directed the dCas9-VPR approximately 1,200 bp upstream of this subfragment(42) (Sup. Table 1). We found that these flies have ectopic sex combs in the T2 and T3 pairs of legs in the first tarsomere, indicating derepression of *Scr* in these discs (Fig. 5B), similar to the flies in which the dCas9-VPR was directed to the *Scr* promoter region (Fig. 1). However, the number of ectopic sex combs in flies where dCas9-VPR was directed to the *Scr*^*E*^ enhancer was significantly lower than that found in flies in which dCas9-VPR was sent to the *Scr* promoter region, as well as the penetrance of the phenotype (Fig. 5B-D). Intriguingly, in the T1 legs, sex combs are also present in the second and even in the third tarsomeres (Fig. 5B-D). Additionally, in some of the T2 legs, we found the same phenotype (Fig. 5C, D). In addition, the average number of sex combs in T1 flies was higher than that in wt flies (Fig. 5C, D), suggesting an enhancement of the expression of *Scr* mediated by the action of dCas9-VPR on the *ScrE* enhancer in T1 (Fig. 5C, D).

**Fig. 5.**
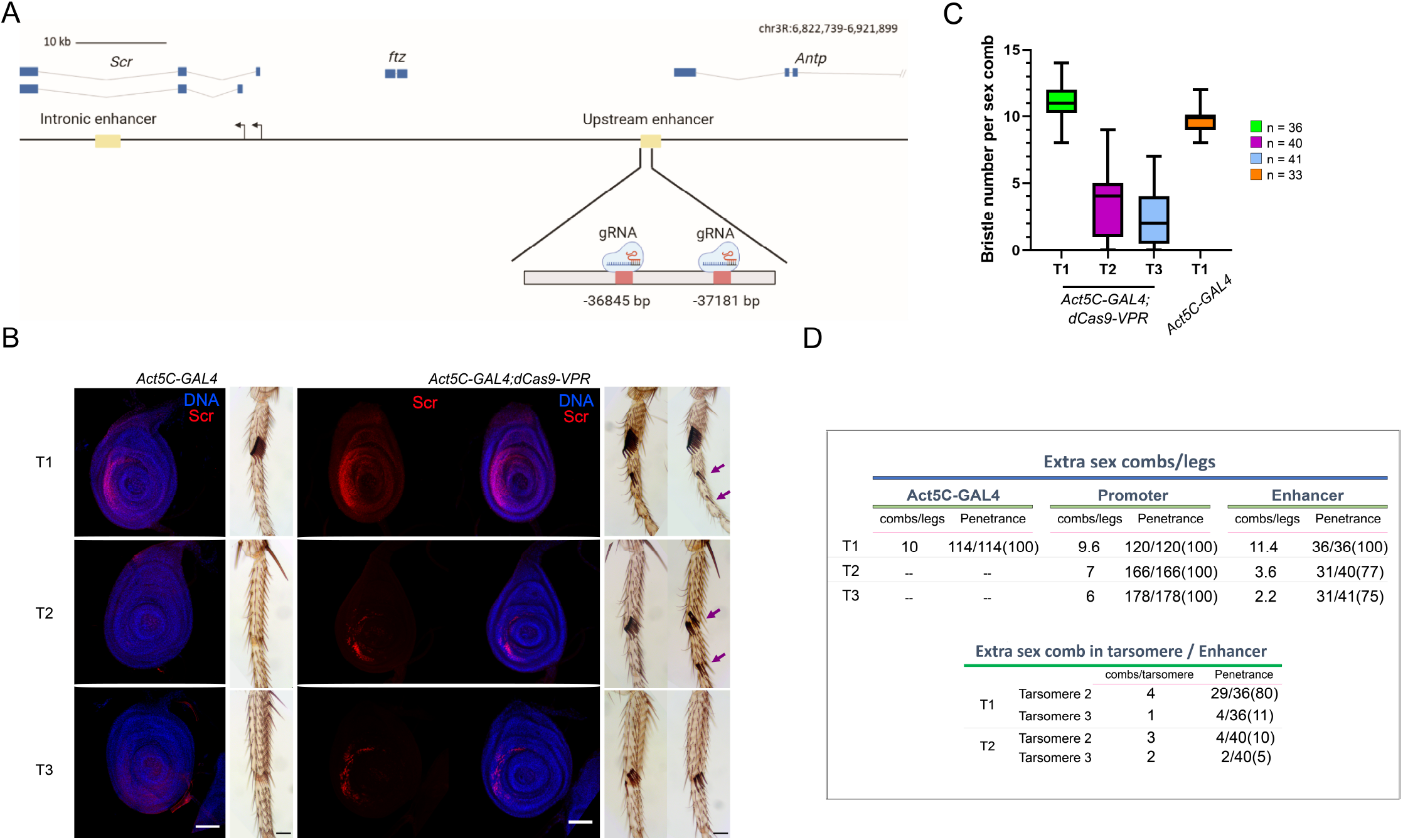
The occupancy of dCas9-VPR in the *Scr* upstream enhancer cause its ectopic expression and the generation of extra sex combs in the 2^nd^ and 3^rd^ pair of legs, as well as in tarsomeres 2 and 3. A) Map of the *Scr* gene showing its genomic organization, the position of enhancers that regulate *Scr*, as well as the position where the gRNAs were directed in the upstream enhancer. The genome coordinates of the gRNAs positions are indicated. B) Extra sex combs phenotypes in male adult legs and the ectopic expression of Scr by immunostaining in leg discs. The arrows indicate the presence of extra sex combs in the tarsomeres (Ts) 2 and 3 in T1 and T2. Barr: 50 μm. C) Quantification of the expressivity indicated by the number of extra sex combs in the legs two and three (T2 and T3) compared with *Act5C*-GAL4 control flies. n=the number of legs analyzed. D) Comparative table between the homeotic transformation phenotype between organisms in which the dCas9-VPR was sent the promoter region versus when it was sent to the upstream enhancer. The penetrance and the expressivity of the phenotypes are indicated.

We followed *Scr* expression through Scr immunostaining of leg discs and found that it was ectopically expressed in T2 and T3 discs. In agreement with the extra sex comb phenotype observed in adult male legs of this genotype, there was a lower number of cells in discs that expressed *Scr* than in discs from flies where dCas9-VPR was directed to the promoter (Fig. 3 vs. Fig. 5). Noticeably, Scr immunostaining was found in a variegated pattern, showing that not all cells responded similarly to *Scr* derepression. Additionally, we observed an increase in the number of cells that expressed *Scr* in the T1 disc, preferentially in the region that will give rise to tarsomeres 2 and 3, explaining the appearance of extra sex combs in these structures (Fig. 5B).

These results show that when dCas9-VPR is sent to the *Scr*-upstream enhancer *Scr*^*E*^, the ectopic expression of *Scr* in the T2 and T3 tarsi occurs at a lower level than when the synthetic transcription factor is sent to the promoter region, but there is also expression outside the disk domain where Scr is normally expressed.

## Discussion

In this work, we analyzed how a synthetic transcription factor, in this case dCas9-VPR, is capable of activating transcription in different types of chromatin. In the case of the subtelomeric element *TAHRE*, cataloged as constitutive heterochromatin and not transcribed in somatic tissues, the artificial activator was very efficient in inducing its transcription from a region corresponding to the 5’ end of *TAHRE*. This is important since it has been reported that nucleosomes can prevent the binding of Cas9 to its target or affect its binding efficiency(56–60). However, the results presented here clearly show that although *TAHRE* is part of the subtelomeric constitutive heterochromatin, dCas9-VPR is capable of binding to its target directed by gRNAs and activating RNPII-mediated transcription. Since dCas9-VPR does not cut DNA and contains three transcription activation domains, it could allow binding to its target in a more stable way even though the chromatin is formed as a nucleosome and, consequently, activate transcription. However, it cannot be ruled out that at some point in the cell cycle, the target DNA could be nucleosome-free, and dCas9-VPR efficiently binds to its target at this point, subsequently inhibiting nucleosome assembly in that region. In any case, dCas9-VPR induces robust transcription of *TAHRE*, suggesting that regions that are configured as constitutive heterochromatin can be transcriptionally activated if a transcription factor has the ability to bind to it.

Regarding the induction of ectopic transcriptional activation of the HOX *Ubx* and *Scr* genes by dCas9-VPR, some points are valuable to discuss. When dCas9-VPR was sent upstream of the *Ubx* promoter, ectopic expression in the wing and eye-antenna discs was variegated. This suggests that the binding of dCas9-VPR in the promoter region was not efficient in causing robust transcriptional activation in all cells or at the same time. Nevertheless, the clonal effect suggests that once *Ubx* is activated in a cell by means of the synthetic transcriptional activator, its expression is maintained in its daughter cells. It is also possible that the genome position to which the transcriptional activator was sent may not be optimal to activate transcription from this promoter, although it has been established that the binding site of the gRNAs approximately -400 bp from the TSS is adequate(26). In any case, in some cells, the expression of *Ubx* was ectopically activated, bypassing the *Ubx* silencing exerted by the Pc group genes in these discs.

On the other hand, ectopic *Scr* activation was more robust (100% penetrance, high expressivity), comparable to typical phenotypes in animals lacking PcG function. As mentioned above, *Scr* has two TSSs, and it is possible that the transcriptional activator would activate both promoters, although this has not been determined. In any case, dCas9-VPR *activated Scr* expression in tissues in which it should be silenced by the PcG genes.

The phenotypes of animals where dCas9-VPR was sent to the *Ubx* or *Scr* enhancers were different from those where it was sent to the promoters. When dCas9-VPR was in the *Scr* upstream enhancer, *Scr* ectopic transcription was activated in T2 and T3 leg discs. However, its expression varied, with low penetrance and expressivity. Noticeably, *Scr* expression was extended toward the center of the disc, mainly in T1, causing the appearance of sexual combs in tarsomeres 2 and 3 of legs derived from these T2 and T3 discs.

The fact that the occupation of the synthetic activator in the *Scr* enhancer is able to activate transcription in the T2 and T3 discs suggests that the enhancer has almost all of the requirements necessary to activate the expression of *Scr* in these tissues and that the addition of a new transcription factor was sufficient to activate CRE. In support of this hypothesis, it has recently been reported in several organisms that the interaction between the enhancer and its target promoter occurs even though the enhancer is not active, both in the tissue where the target gene is expressed and in tissues where it is not(61–63). This observation is relevant from an evolutionary point of view since mutations in Selector genes are known to be the cause of morphological differences between different species. For example, in other *Drosophila* species, the expression of *Scr*, unlike in *D. melanogaster*, causes the presence of sexual combs in tarsomeres 2 and 3(64–66), similar to what we have observed when sending the dCas9-VPR to the *Scr* upstream enhancer.

When dCas9-VPR was sent to the *Ubx bxI* enhancer, it was able to increase *Ubx* transcription in the haltere disc, as determined by measuring the nascent *Ubx* transcript. However, a loss-of-function *Ubx* phenotype is observed in these flies. How is it possible that the effect of locating dCas9-VPR at the *bxI* enhancer is giving results at the same time of both *Ubx* transcriptional activation and repression? This seems to indicate that, as has been previously reported, the occupation of the dCas9-VPR downstream of the TSS, in this case at the intronic *bxI* enhancer, inhibits the passage of RNPII, causing a net effect of *Ubx* silencing. On the other hand, when dCas9-VPR occupies the *bxI* enhancer, it favors its interaction with the *Ubx* promoter, increasing the levels of transcriptional initiation. However, when RNPII reaches the downstream site where dCas9-VPR is positioned, it is stalled and aborted. Nevertheless, we do not know whether these two processes are occurring simultaneously or if the barrier action of dCas9-VPR on RNPII only occurs when the enhancer does not interact with the promoter. In support of this possibility, it has been reported that as transcription increases due to the action of activators in an enhancer, the distance between the promoter and the enhancer increases(67). This could also explain why we observed an *Ubx* variegated expression pattern in haltere discs.

In conclusion, by using the synthetic transcription activator dCas9-VPR, we have shown that it is capable of activating transcription by RNPII, both in regions classified as constitutive (the TAHRE element) and facultative (the HOX genes) heterochromatin regardless of their silenced state. Likewise, the effect on transcriptional activation by sending this artificial activator to a promoter or to an enhancer of the same gene can generate different transcriptional responses in terms of the strength of an ectopic phenotype, as was the case for *Scr*. Surprisingly, the presence of dCas9-VPR in an enhancer within the body of *Ubx* can simultaneously increase the degree of transcription initiation and block the passage of RNPsII in the same gene when located 30 kb from the TSS. All of these findings show that since transcription is a stochastic and plastic mechanism, synthetic transcriptional activators in complex organisms are incorporated in a specific manner into each gene or regulatory region.

## Materials and methods

### gRNAs cloning

Double guides, directed to the different targets of the *Ubx* and *Scr* genes and the retroelement *TAHRE*, were cloned into pCFD4-U6:1-U6:3 (Addgene® plasmid #49411. Tandem expression vector) as previously reported(68). For Gibson Assembly protocol, we used Gibson Assembly® Master Mix de New England BioLabs. All guide sequences were designed using the Benchling platform(69) and Breaking-Cas tool(70). Guide sequences are available in supplementary table 1.

### Transgenic flies

The double sgRNA-plasmids in pCFD4 were integrated into the fly genome at the attP40 integration site on the second chromosome with the phiC31 transformation method by the company BestGene Inc®.

Flies with genotype *sgRNA/CyO;MKRS,Sb/TM6B,Tb,Hu* with each sgRNA sequences directed to *TAHRE, Ubx* or *Scr* or their corresponding enhancers, were crossed with flies *W;If/CyO,UAS:dCas9-VPR/TM6B,Tb,Hu* (donated by Dr. Norbert Perrimon) in order to found the families with genotype *sgRNA/CyO;UAS:dCas9-VPR/TM6B,Tb,Hu*. Finally, we crossed these flies with the corresponding GAL4 driver, accordingly to the experiment. We used *Act5C*-GAL4 driver (Bloomington Stock Center, #4414) and *Tub*-GAL4 (Bloomington Stock Center, #5138) for ubiquitous expression in *TAHRE, Scr* and *Ubx* lines respectively, and specially with *Ubx* promoter lines: *MS1096-GAL4/y*, and *ap>GFP-GAL4/CyO,Tb;+/+* in order to drive expression to the haltere imaginal disc. Fly crosses were conducted at 25°C, unless for *bxI* line, which were performed at 28/18 °C each 12 hours.

### Cuticle treatment

All structures dissected, including antennae, halteres, legs, and wings, were collected from adult fly which were boiled with KOH solution (10% v/v) in a water bath for 5 min. Then, were again boiled in sterile distilled water to remove KOH during 5 min. The structures were dissected in ethanol and mounted in glycerol 50%.

All cuticles dissected had the genotype: *sgRNA/Act5C-GAL4;UAS:dCas9-VPR/+* with the sg-RNA directed to *Ubx* and *Scr* promoters and enhancers as it may apply. As control phenotype we used the corresponding sister lines product of the cross with *Act5C*-GAL4 driver, which genotypes were *sgRNA/CyO;UAS:dCas9-VPR/+* or *Act5C-GAL4/CyO;UAS:dCAs9-VPR/+*.

All cuticles were documented in a Nikon Eclipse E600 upright microscope with a digital camera with Aptina CMOS sensor 5.1 MP, KPA.

### Immunohistochemistry

Wandering-stage third instar larval were dissected in cold PBS 1X and fixed with 4% paraformaldehyde in PBS at room temperature for 30 min. Subsequently, larvae were washed 3 times with PBST (1X PBS and 0.2% Triton) during 15 min each wash. Then, larvae were blocked with BBT (1X PBS, 0.1% BSA and 250 mM NaCl) at 4°C for 1h. Afterwards, the sample was incubated with the primary antibody overnight. After that the larvae were washed again 3 times for 20 min with PBST, the secondary antibody and DAPI was added, and incubated for 2 h at room temperature, and washed 3 times with PBST for 20 min. Finally, the PBST was removed to add the mounting medium and thus extract the imaginal discs for later observation.

In the case of haltere, wing, and eye-antennae imaginal discs for *Ubx* promoter experiments, we analyzed flies with genotype *Ubx-sgRNA/Act5C-GAL4;UAS:dCas9-VPR/+* and lines *Ubx-sgRNA/CyO;UAS:dCas9-VPR/+* or *Act5C-GAL4/CyO;UAS:dCAs9-VPR/+* as siblings lines control. In the case of *bxI* enhancer experiments, we used the haltere imaginal discs with the genotype *bxI-sgRNA/CyO;UAS:dCas9-VPR/αTub-GAL4* for CRISPRa fenotype and *bxI-sgRNA/CyO;UAS:dCas9-VPR/TM6B,Tb,Hu* as the control. Finally, for both *Scr* experiments, promotor (*Scr-*sgRNA) and enhancer (*ScrE-sgRNA*), we performed the immunohistochemistry of leg imaginal discs with genotype *sgRNA/Act5C-GAL4;UAS:dCas9-VPR/+* and lines s*gRNA/CyO;UAS:dCas9-VPR/+* or *Act5C-GAL4/CyO;UAS:dCAs9-VPR/+* as siblings lines control.

Antibodies used were mouse monoclonal antibody Ubx (Ubx FP3.38 DSHB; 1:50), mouse monoclonal antibody Scr (anti-Scr 6H4.1 DSHB; 1:50). As secondary antibody, we used Goat anti-mouse IgG Alexa Fluor 568 (A-11004; Invitrogen) at 1:200 and DAPI concentration 1ng/µL.

Samples were imaged on Olympus FV1000 Multi-photonic Inverted or Olympus FV1000 Confocal Upright microscopes, equipped with a UPLSAPO objective 20X NA: 0.75 AND 60X NA: 1.1.

### Quantitative PCR assay

Adult specimens with genotype *sgRNA/Act5C-GAL4;UAS:dCas9-VPR/+* expressing sgRNAs for TAHRE as CRISPRa lines and the resulting siblings lines from the cross as control were used. Haltere and wing imaginal discs for bxI fly lines with the genotype *bxI-sgRNA/CyO;UAS:dCas9-VPR/αTub-GAL4* were used. The corresponding siblings lines obtained from the cross were used as controls.

The RNA was extracted and purified with TRIzol® Reagent following the manufacturer’s protocol. The isolated RNA was used for the synthesis of cDNA, following the protocol for the M-MLV Reverse Transcriptase enzyme (Invitrogen). End-point PCR amplification of the cDNA was carried out to determine the presence of TAHRE transcripts using Taq polymerase. RP49 was used as a reference gene.

The qRT-PCR experiments were conducted using LightCycler® FastStart DNA Master PLUS SYBR Green I (Roche) for TAHRE experiments and Maxima SYBR Green/ROX qPCR Master Mix (Thermo Scientific™) for the bxI enhancer experiments in a Roche Real Time PCR LightCycler® 1.5. The threshold cycle (C_T_ or 2^-ΔΔCT^) method(71) was used to calculate the fold-change for transcript relative quantification. At least three independent biological replicates were analyzed in each case. Statistical analyzes (student t and One-way ANOVA tests) were performed using the GraphPad Prism 8 software. The primers used for qRT-PCR experiments for each target are listed in suppl. Table 2.

### Public ATAC-seq data analysis

ATAC-seq data from *Drosophila melanogaster* haltere were previously published(48). Raw data was downloaded from the GEO database (GSE166714). Fastq files were aligned to the dm6 genome using Bowtie2 software and peak calling was performed using MACS3. BigWig files were processed using Deeptools library and finally were visualized using the IGV software.

## Acknowledgments

We thank Dr. Robert Perrimon for the donation of the *UAS:dCas9-VPR* fly. We also thank Arturo Pimentel, Andres Saralegui, Chris Wood and the National Laboratory of Advance Microscopy at the Instituto de Biotecnología/UNAM for advice in the use of the confocal microscopes. This work was supported by the grants: PAPIIT-UNAM No. IN200218 and CONACyT infraestructura No. 316070 AOY and SCR are supported by CONACyT scholarships

## Supplementary Figures legends

**Sup. Fig. 1.**
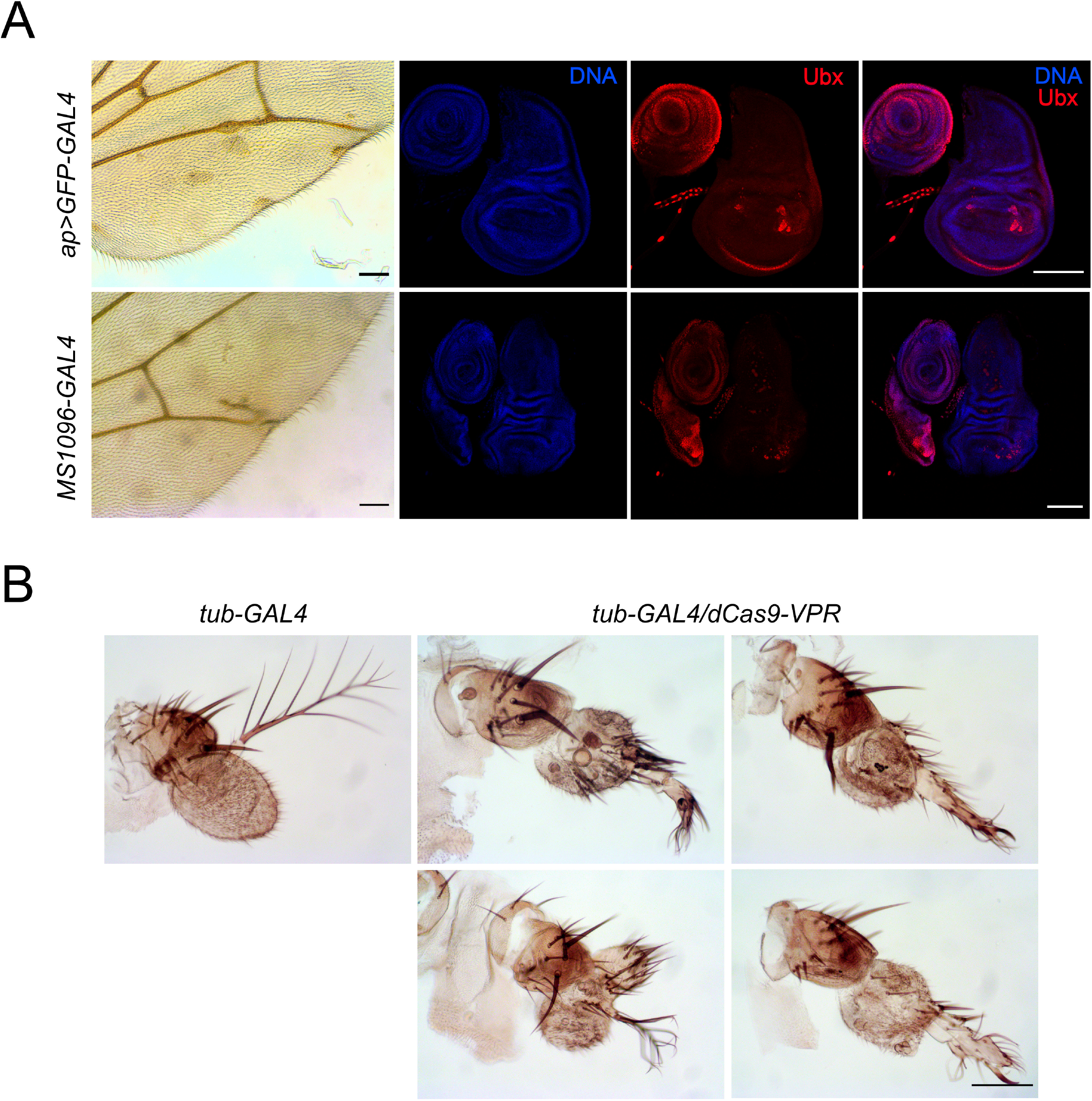
*Ubx* ectopic expression by the Cas9-VPR directed to its promoter using different GAL4-drivers. A) Expression of *Ubx* in the wing disc modulated by the wing specific drivers *apterus* and MS1096. Note that the *Ubx* expression preferentially occurs in the posterior region of the disc in both cases. B) Transformation of antennas to legs using the *tubulin*-GAL4 driver. Note that the expressivity of the phenotype is more dramatic than when the *Actin5*-driver is used. The corresponding genotypes are indicated in the figure.

**Sup. Fig. 2.**
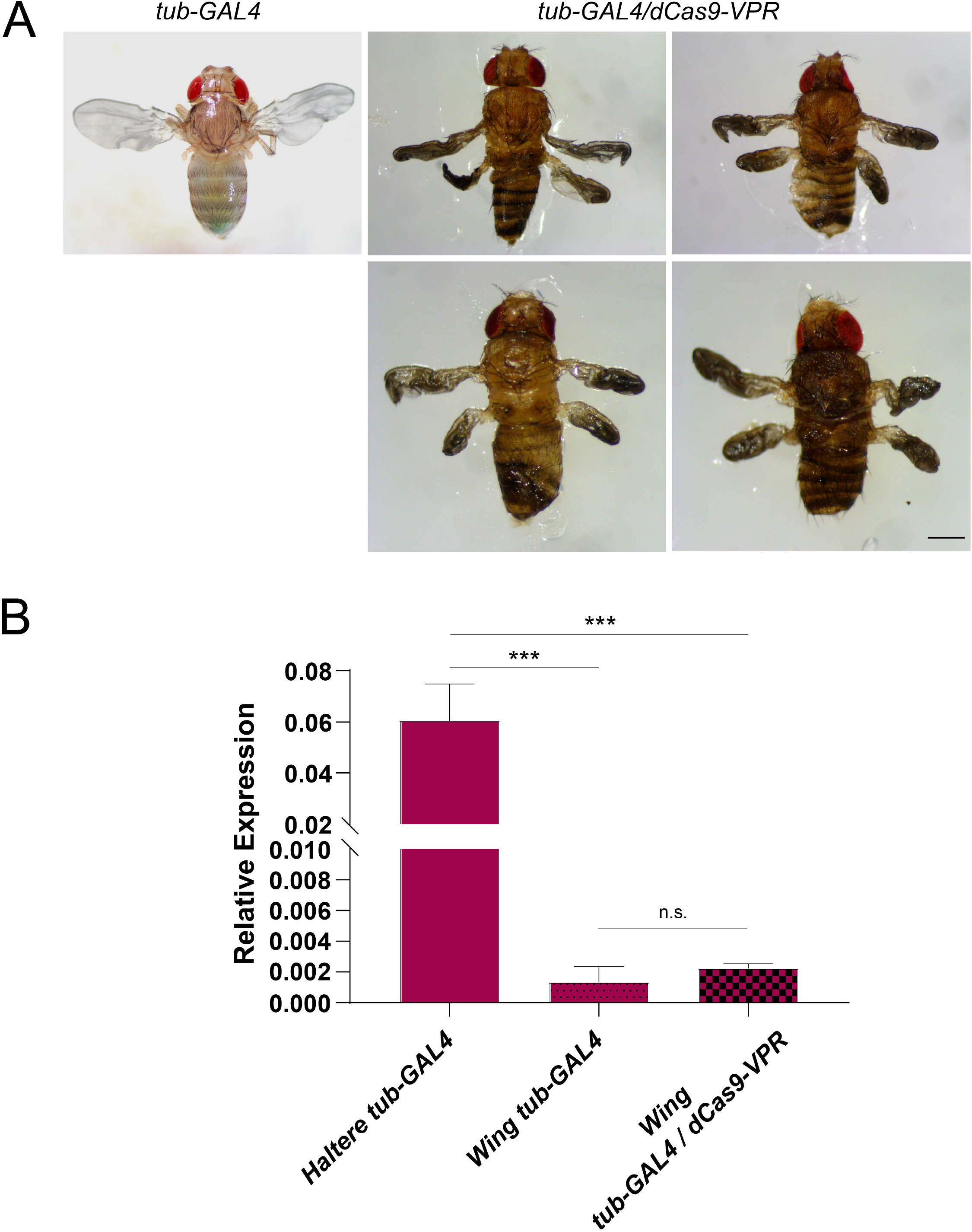
Loss of function phenotypes and quantification of the expression of *Ubx* in wing discs by sending the dCas9 to the *bxI* enhancer using the tubulin driver. A) Examples of pharates that cannot hatch when dCas9-VPR is sent to the *bxI* enhancer using the tubulin-GAL4 driver, showing almost complete transformation of haleteres to wings. B). Quantitative qRT-PCR analysis of the *Ubx* nascent transcript in wing discs of flies in which dCas9-VPR is sent to the *bxI* enhancer. Note that that unlike the case of the haltere disc, the Cas9-VPR does not enhance the *Ubx* transcription in this tissue.

## Supplementary Tables

**Table 1.**
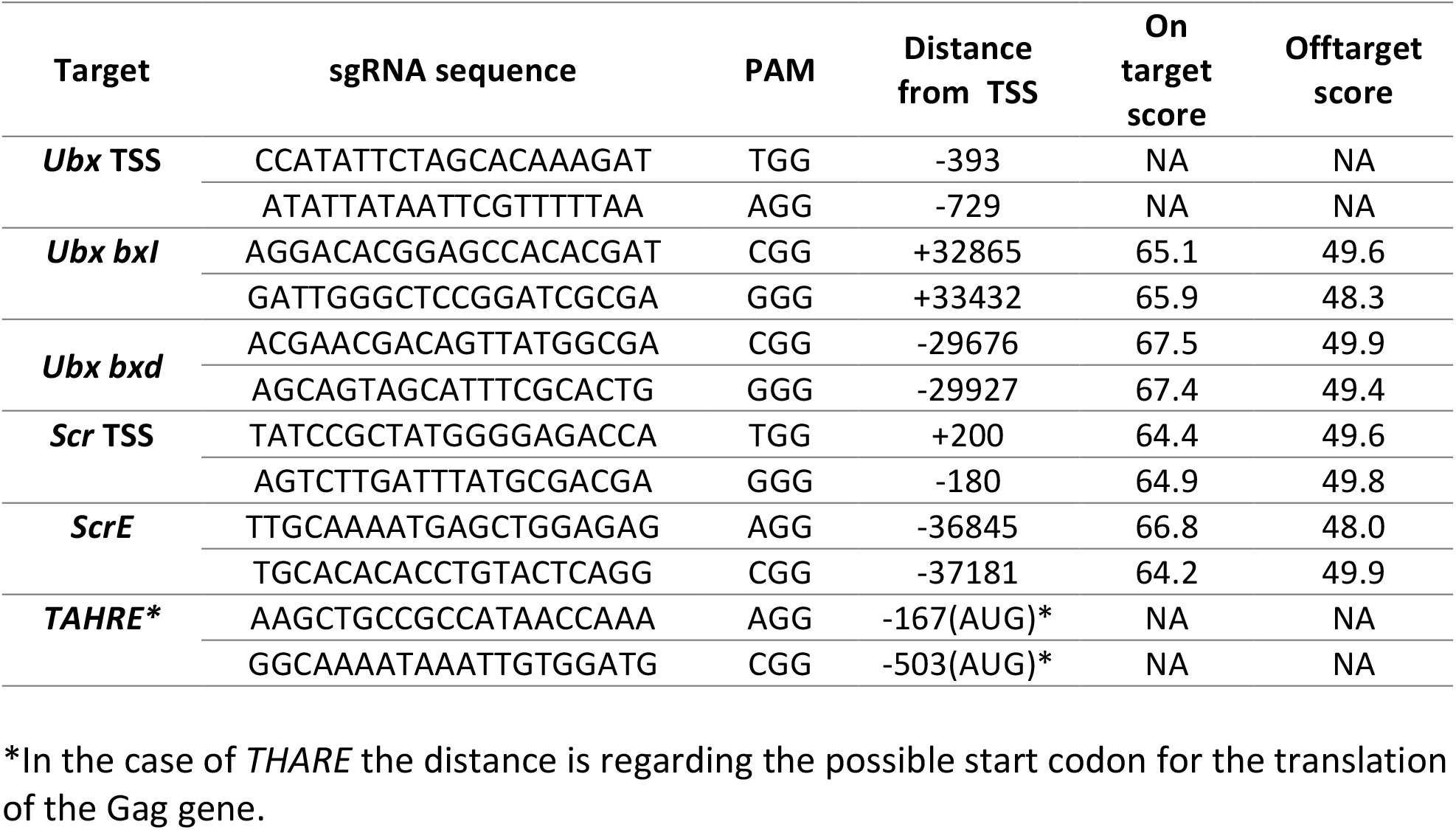
Sequence of the different gRNA’s used in this work.

**Table 2.**
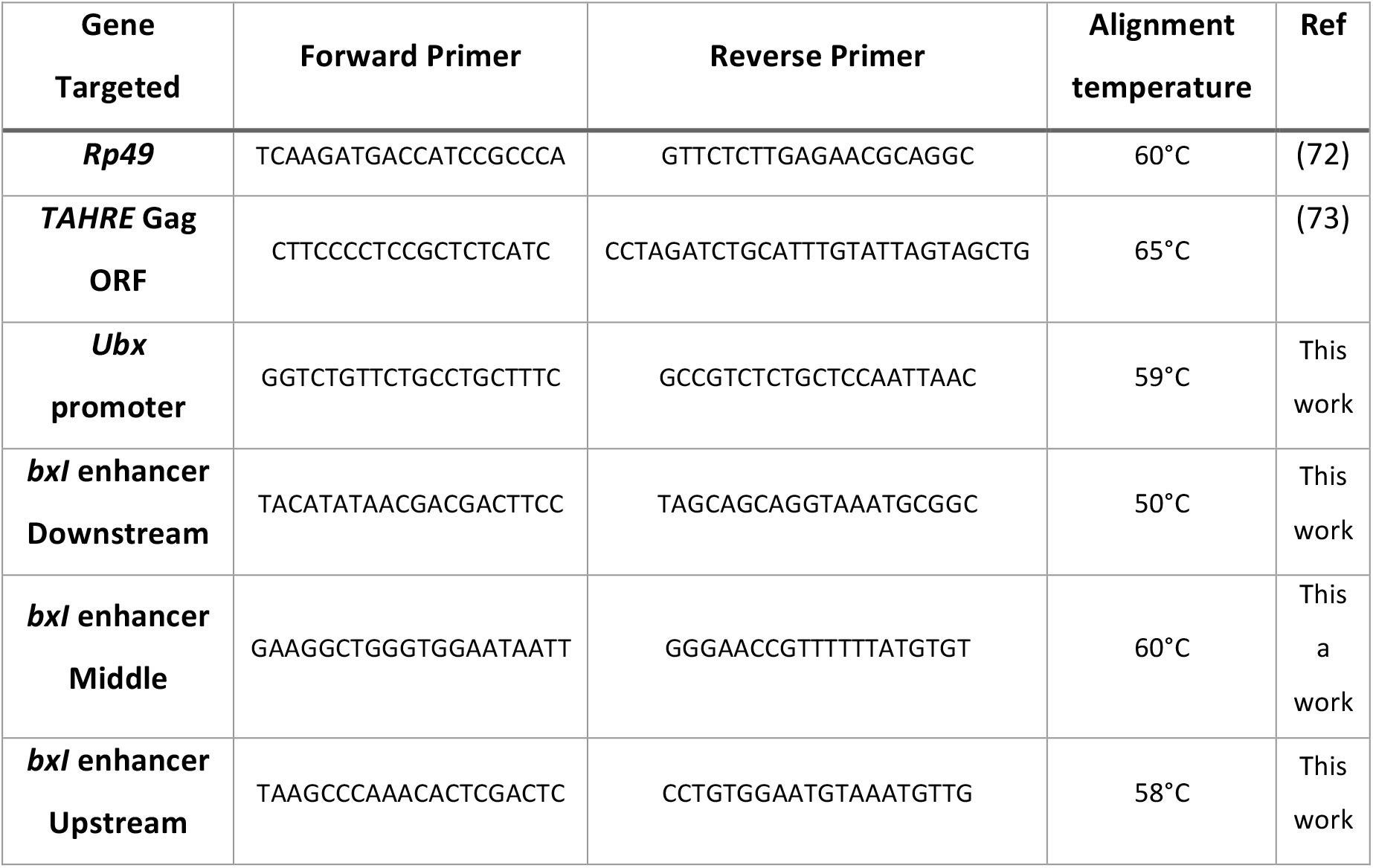
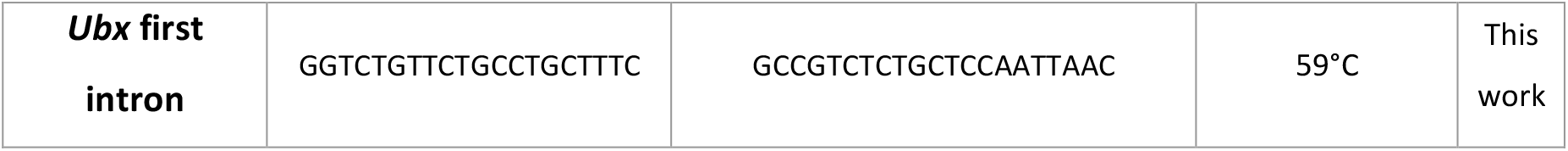
Primers used for RT-qPCR for each region analyzed

**Table 3.** Viability and penetrance for homeotic transformations in *Ubx TSS* fly lines.

|  | Viability | Homeotic transformation |
| --- | --- | --- |
| ***Ubx-sgRNA/Act5C-GAL4;UAS:dCas9-VPR/+*** | 59/59 (100) | 37/59 (63%) |
| *sgRNA/Act5C-GAL4;MKRS/+* | 58/59 (98) | 0 |
| *Act5C-GAL4/CyO;UAS:dCas9-VPR/+* | 53/59 (90) | 0 |
| *sgRNA/CyO; UAS:dCas9-VPR/+* | 53/59 (90) | 0 |
| *Act5C-GAL4/CyO;MKRS/+* | 57/59 (97) | 0 |
| *sgRNA/CyO; MKRS/+* | 57/59 (97) | 0 |

**Table 4.**
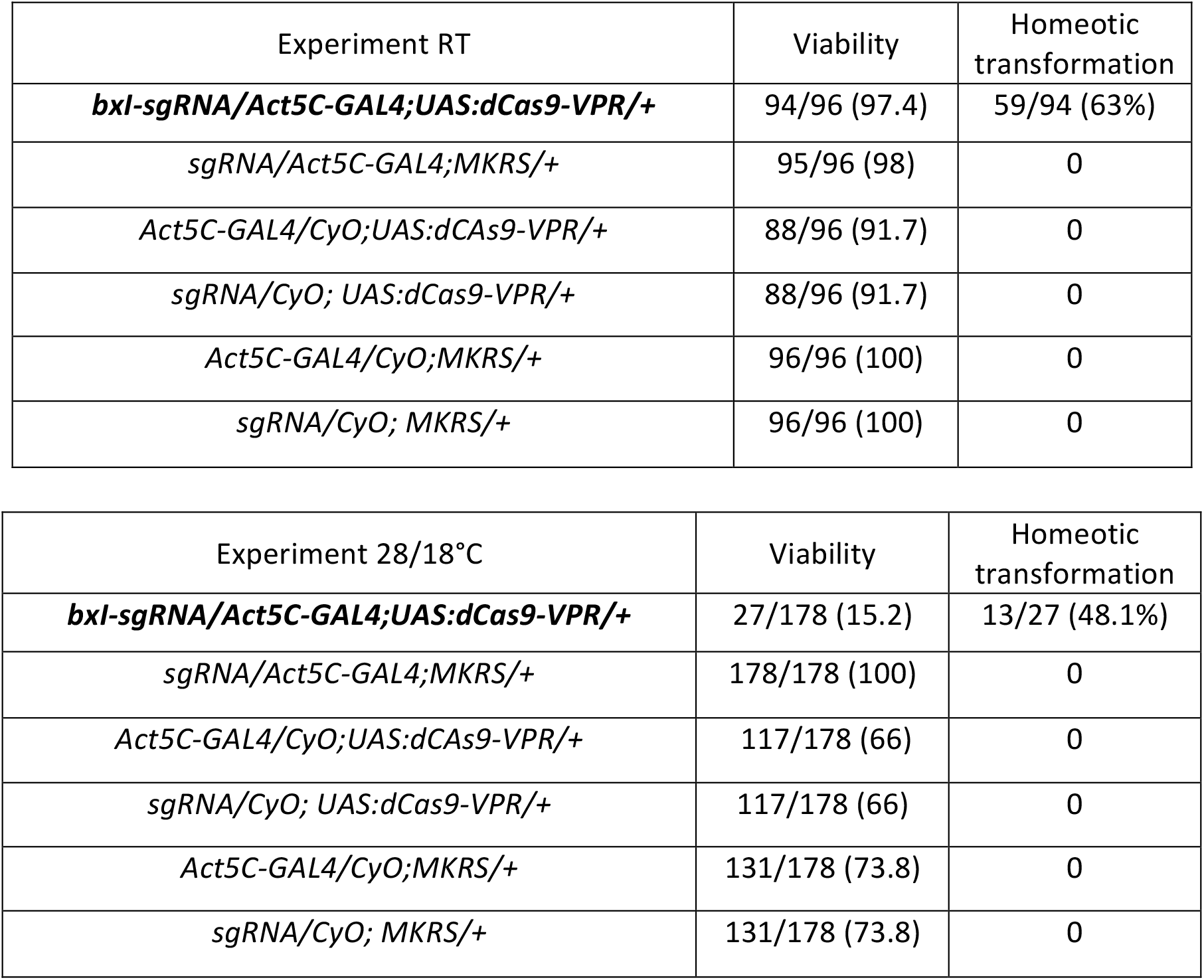
Viability and penetrance for homeotic transformations in *Ubx bxI* organisms.

| Experiment RT | Viability | Homeotic transformation |
| --- | --- | --- |
| ***bxl-sgRNA/Act5C-GAL4;UAS:dCas9-VPR/+*** | 94/96 (97.4) | 59/94 (63%) |
| *sgRNA/Act5C-GAL4;MKRS/+* | 95/96 (98) | 0 |
| *Act5C-GAL4/CyO;UAS:dCas9-VPR/+* | 88/96 (91.7) | 0 |
| *sgRNA/CyO; UAS:dCas9-VPR/+* | 88/96 (91.7) | 0 |
| *Act5C-GAL4/CyO;MKRS/+* | 96/96 (100) | 0 |
| *sgRNA/CyO; MKRS/+* | 96/96 (100) | 0 |

| Experiment 28/18°C | Viability | Homeotic transformation |
| --- | --- | --- |
| ***bxl-sgRNA/Act5C-GAL4;UAS:dCas9-VPR/+*** | 27/178 (15.2) | 13/27 (48.1%) |
| *sgRNA/Act5C-GAL4;MKRS/+* | 178/178 (100) | 0 |
| *Act5C-GAL4/CyO;UAS:dCas9-VPR/+* | 117/178 (66) | 0 |
| *sgRNA/CyO; UAS:dCas9-VPR/+* | 117/178 (66) | 0 |
| *Act5C-GAL4/CyO;MKRS/+* | 131/178 (73.8) | 0 |
| *sgRNA/CyO; MKRS/+* | 131/178 (73.8) | 0 |

## References

1. H. Chen, B. F. Pugh, What do Transcription Factors Interact With? J. Mol. Biol. 433, 166883 (2021).

2. S. Schilbach, et al., Structures of transcription pre-initiation complex with TFIIH and Mediator. Nature 551, 204–209 (2017).

3. J. M. Tome, N. D. Tippens, J. T. Lis, Single-molecule nascent RNA sequencing identifies regulatory domain architecture at promoters and enhancers. Nat. Genet. 50, 1533–1541 (2018).

4. F. Le Dily, M. Beato, Signaling by Steroid Hormones in the 3D Nuclear Space. Int. J. Mol. Sci. 19 (2018).

5. J. Judd, F. M. Duarte, J. T. Lis, Pioneer-like factor GAF cooperates with PBAP (SWI/SNF) and NURF (ISWI) to regulate transcription. Genes Dev. 35, 147–156 (2021).

6. S. Heinz, C. E. Romanoski, C. Benner, C. K. Glass, The selection and function of cell type-specific enhancers (2015) https://doi.org/10.1038/nrm3949 (February 8, 2022).

7. A. Field, K. Adelman, Evaluating Enhancer Function and Transcription. Annu. Rev. Biochem. 89, 213–234 (2020).

8. M. S. C. Larke, et al., Enhancers predominantly regulate gene expression during differentiation via transcription initiation. Mol. Cell 81, 983-997.e7 (2021).

9. B. Lim, M. S. Levine, Enhancer-promoter communication: hubs or loops? Curr. Opin. Genet. Dev. 67, 5–9 (2021).

10. A. Gurumurthy, Y. Shen, E. M. Gunn, J. Bungert, Phase Separation and Transcription Regulation: Are Super-Enhancers and Locus Control Regions Primary Sites of Transcription Complex Assembly? Bioessays 41 (2019).

11. A. Boija, et al., Transcription Factors Activate Genes through the Phase-Separation Capacity of Their Activation Domains. Cell 175, 1842-1855.e16 (2018).

12. L. Hilbert, et al., Transcription organizes euchromatin via microphase separation. Nat. Commun. 12 (2021).

13. D. Chetverina, M. Erokhin, P. Schedl, GAGA factor: a multifunctional pioneering chromatin protein. Cell. Mol. Life Sci. 78, 4125–4141 (2021).

14. K. S. Zaret, Pioneer Transcription Factors Initiating Gene Network Changes. Annu. Rev. Genet. 54, 367–385 (2020).

15. R. M. Marsano, E. Giordano, G. Messina, P. Dimitri, A New Portrait of Constitutive Heterochromatin: Lessons from Drosophila melanogaster. Trends Genet. 35, 615–631 (2019).

16. Y. Guo, S. Zhao, G. G. Wang, Polycomb Gene Silencing Mechanisms: PRC2 Chromatin Targeting, H3K27me3 “Readout”, and Phase Separation-Based Compaction. Trends Genet. 37, 547–565 (2021).

17. G. van Mierlo, G. J. C. Veenstra, M. Vermeulen, H. Marks, The Complexity of PRC2 Subcomplexes. Trends Cell Biol. 29, 660–671 (2019).

18. M. I. Kuroda, H. Kang, S. De, J. A. Kassis, Dynamic Competition of Polycomb and Trithorax in Transcriptional Programming. Annu. Rev. Biochem. 89, 235–253 (2020).

19. S. Z. Deluca, M. Ghildiyal, L. Y. Pang, A. C. Spradling, Differentiating Drosophila female germ cells initiate Polycomb silencing by regulating PRC2-interacting proteins. Elife 9, 1–33 (2020).

20. J. Liu, M. Ali, Q. Zhou, Establishment and evolution of heterochromatin. Ann. N. Y. Acad. Sci. 1476, 59–77 (2020).

21. A. Martella, et al., Systematic Evaluation of CRISPRa and CRISPRi Modalities Enables Development of a Multiplexed, Orthogonal Gene Activation and Repression System. ACS Synth. Biol. 8, 1998–2006 (2019).

22. A. Chavez, et al., Highly efficient Cas9-mediated transcriptional programming. Nat. Methods 12, 326–328 (2015).

23. S. Lin, B. Ewen-Campen, X. Ni, B. E. Housden, N. Perrimon, In vivo transcriptional activation using CRISPR/Cas9 in Drosophila. Genetics 201, 433–442 (2015).

24. N. Matharu, et al., CRISPR-mediated activation of a promoter or enhancer rescues obesity caused by haploinsufficiency. Science (80-.). 363 (2019).

25. J. Fitz, et al., Spt5-mediated enhancer transcription directly couples enhancer activation with physical promoter interaction. Nat. Genet. 52, 505–515 (2020).

26. B. Ewen-Campen, et al., Optimized strategy for in vivo Cas9-activation in Drosophila. Proc. Natl. Acad. Sci. U. S. A. 114, 9409–9414 (2017).

27. J. A. Kassis, J. A. Kennison, J. W. Tamkun, Polycomb and Trithorax Group Genes in Drosophila. Genetics 206, 1699–1725 (2017).

28. D. N. Markova, S. M. Christensen, E. Betrán, Telomere-Specialized Retroelements in Drosophila: Adaptive Symbionts of the Genome, Neutral, or in Conflict? Bioessays 42 (2020).

29. S. Cacchione, G. Cenci, G. D. Raffa, Silence at the End: How Drosophila Regulates Expression and Transposition of Telomeric Retroelements. J. Mol. Biol. 432, 4305–4321 (2020).

30. A. Penagos-Puig, M. Furlan-Magaril, Heterochromatin as an Important Driver of Genome Organization. Front. cell Dev. Biol. 8 (2020).

31. J. M. Gibert, F. Peronnet, The Paramount Role of Drosophila melanogaster in the Study of Epigenetics: From Simple Phenotypes to Molecular Dissection and Higher-Order Genome Organization. Insects 12 (2021).

32. P. K. Rivlin, A. Gong, A. M. Schneiderman, R. Booker, The role of Ultrabithorax in the patterning of adult thoracic muscles in Drosophila melanogaster. Dev. Genes Evol. 2001 2112 211, 55–66 (2001).

33. V. Pirrotta, Chi Shing Chan, D. McCabe, S. Qian, Distinct parasegmental and imaginal enhancers and the establishment of the expression pattern of the Ubx gene. Genetics 141, 1439–1450 (1995).

34. S. M. Smolik-Utlaut, Dosage requirements of Ultrabithorax and bithoraxoid in the determination of segment identity in Drosophila melanogaster. Genetics 124, 357–366 (1990).

35. T. Fuqua, et al., Dense and pleiotropic regulatory information in a developmental enhancer. Nature 587, 235–239 (2020).

36. R. K. Delker, V. Ranade, R. Loker, R. Voutev, R. S. Mann, Low affinity binding sites in an activating CRM mediate negative autoregulation of the Drosophila Hox gene Ultrabithorax. PLoS Genet. 15 (2019).

37. R. Cao, et al., Role of histone H3 lysine 27 methylation in Polycomb-group silencing. Science 298, 1039–1043 (2002).

38. R. K. Maeda, F. Karch, The bithorax complex of Drosophila an exceptional Hox cluster. Curr. Top. Dev. Biol. 88, 1–33 (2009).

39. L. C. Yao, G. J. Liaw, C. Y. Pai, Y. H. Sun, A common mechanism for antenna-to-Leg transformation in Drosophila: suppression of homothorax transcription by four HOM-C genes. Dev. Biol. 211, 268–276 (1999).

40. D. L. Brower, Ultrabithorax gene expression in Drosophila imaginal discs and larval nervous system. Development 101, 83–92 (1987).

41. R. A. H. White, M. Wilcox, Distribution of Ultrabithorax proteins in Drosophila. EMBO J. 4, 2035–2043 (1985).

42. S. E. Eksi, O. Barmina, C. L. McCallough, A. Kopp, T. V. Orenic, A Distalless-responsive enhancer of the Hox gene Sex combs reduced is required for segment-and sex-specific sensory organ development in Drosophila. PLoS Genet. 14, e1007320 (2018).

43. J. W. Southworth, J. A. Kennison, Transvection and Silencing of the Scr Homeotic Gene of Drosophila melanogaster. Genetics 161, 733–746 (2002).

44. J. M. Calvo, P. Carmen, Sex combs reduced (Scr) regulatory region of Drosophila revisited. Mol. Genet. Genomics 292, 773–787 (2017).

45. M. O. Starr, et al., Molecular dissection of cis-regulatory modules at the Drosophila bithorax complex reveals critical transcription factor signature motifs. Dev. Biol. 359, 290–302 (2011).

46. S. Qian, M. Capovilla, V. Pirrotta, Molecular mechanisms of pattern formation by the BRE enhancer of the Ubx gene. EMBO J. 12, 3865–3877 (1993).

47. S. Qian, M. Capovilla, V. Pirrotta, The bx region enhancer, a distant cis-control element of the Drosophila Ubx gene and its regulation by hunchback and other segmentation genes. EMBO J. 10, 1415–25 (1991).

48. R. Loker, J. E. Sanner, R. S. Mann, Cell-type-specific Hox regulatory strategies orchestrate tissue identity. Curr. Biol. 31, 4246-4255.e4 (2021).

49. L. Sipos, G. Kozma, E. Molnár, W. Bender, In situ dissection of a Polycomb response element in Drosophila melanogaster. Proc. Natl. Acad. Sci. U. S. A. 104, 12416–12421 (2007).

50. J. Zirin, J. Bosch, R. Viswanatha, S. E. Mohr, N. Perrimon, State-of-the-art CRISPR for in vivo and cell-based studies in Drosophila. Trends Genet. (2021) https://doi.org/10.1016/J.TIG.2021.11.006 (January 29, 2022).

51. L. A. Gilbert, et al., CRISPR-mediated modular RNA-guided regulation of transcription in eukaryotes. Cell 154, 442 (2013).

52. L. S. Qi, et al., Repurposing CRISPR as an RNA-guided platform for sequence-specific control of gene expression. Cell 152, 1173–1183 (2013).

53. Y. Zhu, et al., Predicting enhancer transcription and activity from chromatin modifications. Nucleic Acids Res. 41, 10032–10043 (2013).

54. O. Mikhaylichenko, et al., The degree of enhancer or promoter activity is reflected by the levels and directionality of eRNA transcription. Genes Dev. 32, 42–57 (2018).

55. F. Liu, Enhancer-derived RNA: A Primer. Genomics. Proteomics Bioinformatics 15, 196–200 (2017).

56. M. A. Horlbeck, et al., Nucleosomes impede cas9 access to DNA in vivo and in vitro. Elife 5, 1–21 (2016).

57. R. S. Isaac, et al., Nucleosome breathing and remodeling constrain CRISPR-Cas9 function. Elife 5, 1–14 (2016).

58. K. T. Jensen, et al., Chromatin accessibility and guide sequence secondary structure affect CRISPR-Cas9 gene editing efficiency. FEBS Lett. 591, 1892–1901 (2017).

59. R. M. Yarrington, S. Verma, S. Schwartz, J. K. Trautman, D. Carroll, Nucleosomes inhibit target cleavage by CRISPR-Cas9 in vivo. Proc. Natl. Acad. Sci. U. S. A. 115, 9351–9358 (2018).

60. S. A. Verkuijl, M. G. Rots, The influence of eukaryotic chromatin state on CRISPR-Cas9 editing efficiencies. Curr. Opin. Biotechnol. 55, 68–73 (2019).

61. O. Kyrchanova, P. Georgiev, Mechanisms of Enhancer-Promoter Interactions in Higher Eukaryotes. Int. J. Mol. Sci. 2021, Vol. 22, Page 671 22, 671 (2021).

62. Y. Ghavi-Helm, et al., Enhancer loops appear stable during development and are associated with paused polymerase. Nature 512, 96–100 (2014).

63. A. J. Rubin, et al., Lineage-specific dynamic and pre-established enhancer– promoter contacts cooperate in terminal differentiation. Nat. Genet. 2017 4910 49, 1522–1528 (2017).

64. K. Tanaka, O. Barmina, L. E. Sanders, M. N. Arbeitman, A. Kopp, Evolution of sex-specific traits through changes in HOX-dependent doublesex expression. PLoS Biol. 9, e1001131 (2011).

65. O. Barmina, A. Kopp, Sex-specific expression of a HOX gene associated with rapid morphological evolution. 311, 277–286 (2007).

66. J. Atallah, H. Watabe, A. Kopp, Many ways to make a novel structure: a new mode of sex comb development in Drosophilidae. Evol. Dev. 14, 476–483 (2012).

67. N. S. Benabdallah, et al., Decreased Enhancer-Promoter Proximity Accompanying Enhancer Activation. Mol. Cell 76, 473-484.e7 (2019).

68. F. Port, H. M. H.-M. Chen, T. Lee, S. L. Bullock, Optimized CRISPR/Cas tools for efficient germline and somatic genome engineering in Drosophila. Proc. Natl. Acad. Sci. 111, E2967–E2976 (2014).

69. Benchling, Cloud-based informatics platform for life sciences R&D | Benchling (2021) (January 20, 2022).

70. J. C. Oliveros, et al., Breaking-Cas-interactive design of guide RNAs for CRISPR-Cas experiments for ENSEMBL genomes. Nucleic Acids Res. 44, W267–W271 (2016).

71. T. D. Schmittgen, K. J. Livak, Analyzing real-time PCR data by the comparative CT method. Nat. Protoc. 3, 1101–1108 (2008).

72. Z. Palomera-Sanchez, A. Bucio-Mendez, V. Valadez-Graham, E. Reynaud, M. Zurita, Drosophila p53 Is Required to Increase the Levels of the dKDM4B Demethylase after UV-induced DNA Damage to Demethylate Histone H3 Lysine 9. J. Biol. Chem. 285, 31370–31379 (2010).

73. M. F. Walter, et al., Effects of telomere length in Drosophila melanogaster on life span, fecundity, and fertility. Chromosoma 116, 41–51 (2007).

